# Cerebral Cavernous Malformations Develop through Clonal Expansion of Mutant Endothelial Cells

**DOI:** 10.1101/405191

**Authors:** Matthew R. Detter, Daniel A. Snellings, Douglas A. Marchuk

## Abstract

**Rationale:** Vascular malformations arise in vessels throughout the entire body. Causative genetic mutations have been identified for many of these diseases; however, little is known about the mutant cell lineage within these malformations.

**Objective:** We utilize an inducible mouse model of cerebral cavernous malformations (CCMs) coupled with a multi-color fluorescent reporter to visualize the contribution of mutant endothelial cells (ECs) to the malformation.

**Methods and Results:** We combined a Ccm3 mouse model with the confetti fluorescent reporter to simultaneously delete *Ccm3* and label the mutant EC with one of four possible colors. We acquired Z-series confocal images from serial brain sections and created 3D reconstructions of entire CCMs to visualize mutant ECs during CCM development. We observed a pronounced pattern of CCMs lined with mutant ECs labeled with a single confetti color (n=42). The close 3D distribution, as determined by the nearest neighbor analysis, of the clonally dominant ECs within the CCM was statistically different than the background confetti labeling of ECs in non-CCM control brain slices as well as a computer simulation (p<0.001). Many of the small (<100μm diameter) CCMs consisted, almost exclusively, of the clonally dominant mutant ECs labeled with the same confetti color whereas the large (>100μm diameter) CCMs contained both the clonally dominant mutant cells and wildtype ECs. We propose of model of CCM development in which an EC acquires a second somatic mutation, undergoes clonal expansion to initiate CCM formation, and then incorporates neighboring wildtype ECs to increase the size of the malformation.

**Conclusions:** This is the first study to visualize, with single-cell resolution, the clonal expansion of mutant ECs within CCMs. The incorporation of wildtype ECs into the growing malformation presents another series of cellular events whose elucidation would enhance our understanding of CCMs and may provide novel therapeutic opportunities.

## Introduction

Vascular malformations occur in arterial, capillary, venous, and lymphatic vessels throughout the entire body. Advances in DNA sequencing capabilities have led to an extensive list of pathogenic loss, or gain, -of-function mutations that cause these diseases, as reviewed previously.^1, 2^ While specific mutations have been identified for many of these malformations, what is less understood is how a mutant cell, harboring one of these mutations, leads to malformation development and growth. More specifically, it is not known if a single mutation event is sufficient to form these malformations or if multiple different mutation events are necessary to form these complex and structurally abnormal vessels. In this study we utilize an inducible mouse model to investigate this question of mutant endothelial cell (EC) lineage in a specific vascular disease, cerebral cavernous malformations (CCMs).

Cerebral cavernous malformations are ectatic, capillary-venous vessels which are lined with a single layer of endothelial cells and lack surrounding parenchymal cells. Disruption of the endothelial barrier within these slow-flow malformations leads to recurrent hemorrhages, seizures, and focal neurologic deficits. CCMs develop following bi-allelic inactivating mutations in *KRIT1/CCM1* (*KREV1/RAP1A interaction trapped-1*)^3, 4^, *OSM*/*CCM2* (*Osmosensing scaffold for MEKK3*)^5, 6^, or *PDCD10*/*CCM3* (*Programmed cell death 10*)^7^. CCMs develop sporadically or through an autosomal-dominant pattern of inheritance. Harboring a germline loss-of-function mutation, patients develop the heritable form of disease when a second somatic mutation occurs in the single remaining functional *CCM* allele. The familial disease often results in 10s to 100s of CCMs developing in adolescent patients while the sporadic form of disease typically presents as a solitary CCM in adult patients. This difference in lesion burden is analogous to Knudson’s observations of tumor burden in sporadic and familial forms of pediatric retinoblastoma^8^. Knudson’s two-hit model, of the first mutation in germ cells and the second mutation occurring in somatic cells, has been employed to explain CCM pathogenesis. DNA sequencing studies have identified these second somatic mutations, with low allele frequencies, in human CCM samples.^9-12^ What remains unknown is if a single second somatic mutation is sufficient for CCM formation and precisely where the mutant cells reside within the complex cellular architecture of the vascular malformation. We combined the neonatal Ccm3 mouse model with a multi-color fluorescent reporter to label mutant ECs and directly observe how mutant cells contribute to CCM pathology.

## Methods

### Nomenclature

We have employed the conventional and distinct rules of nomenclature for disease, lesion, human gene, and mouse gene. The human inherited disease Cerebral Cavernous Malformation, or the vascular lesion (in any species) that characterizes the phenotype, is abbreviated as CCM. The human *CCM3* gene is fully capitalized and italicized; the murine *Ccm3* gene is italicized with both upper- and lower-case letters.

### Mouse Procedures

All animal procedures were approved by the Duke University Institutional Animal Care and Use Committee. The endothelial-specific, tamoxifen-inducible cre recombinase (*PDGFb-iCreERT2*)^13^, lox-P flanked and knockout *Ccm3* alleles^14^, and the Brainbow2.1, commonly referred to as confetti, reporter allele (*R26R-Confetti*)^15, 16^ were bred into C57BL/6J mice with the following experimental genotype: *PDGFb-iCreERT2*, *Ccm3*^*fl/KO*^, *R26R-Confetti*^*fl/wt*^. The control C57BL/6J mice had the following control genotype: *PDGFb-iCreERT2*, *Ccm3*^*wt/wt*^, *R26R-Confetti*^*fl/wt*^. A single intragastric injection of 2μg of tamoxifen (Sigma T5648), dissolved in a 9:1 corn oil to ethanol solution, was administered to experimental mice on postnatal day (P) 3, 4, or 5 and on P3 for the control mice. The experimental mice injected on P3 were euthanized on P39, P4 on P48, and P5 on P53. The control mouse injected on P3 was euthanized on P25.

### Brain Processing

Following CO_2_ euthanasia, we performed cardiac perfusion with phosphate buffered saline (PBS) and 4% paraformaldehyde (PFA). The brains and eyes were carefully dissected in stored in 4% PFA overnight at 4C. Serial sectioning of the brains into 80-μm thick slices was performed with a vibrating microtome (Leica VT1000 S, Frequency: 8, Speed: 5). The slices were mounted onto slides with Fluoromount-G (SouthernBiotech 0100-01) for confocal microscopy.

### Retina Immunofluorescence

Heat-induced antigen retrieval of the dissected retinas was performed with 10mM citric acid with 0.05% Tween20 (J.T. Baker X251-07) (pH 6) buffer for 5 minutes at 85C. Retinas were permeabilized with 1.0% triton X-100 (Sigma X100) blocked with 10% normal goat serum (Sigma G6767), incubated with rabbit anti-pMLC2 (1:200, Cell Signaling 3674S) overnight at 4C, incubated with Alexa Fluor 647 goat anti-rabbit IgG (Invitrogen A21244) for 2 hours at room temp, and mounted onto slices with Fluoromount-G (SouthernBiotech 0100-01) for confocal microscopy.

### Confocal Microscopy

Images were acquired with a Zeiss 880 inverted confocal microscope equipped with GaAsP high QE 32 channel spectral array detector and 2 standard alkali PMTs for far-red detection. The brain slices were imaged with a 20x (0.80 NA) air objective. The retinas were imaged with the same 20x (0.8 NA) air objective and a 40x (1.2 NA) water objective. A Z-series for each brain region of interest was acquired through the full thickness of the sample with a step size of 2.5μm.

### 3D Image Reconstruction

Z-series from serial slices, each containing part of the CCM of interest, were aligned, utilizing white and gray matter boundaries, and concatenated in ImageJ. Uniform adjustments of brightness, contrast, and gamma for the concatenated images were performed with IMARIS 9.0. Each channel was assigned its respective confetti color for display. The surface tool of IMARIS was used to manually outline the vascular lumens of the malformations to provide a visual aid in identifying the CCMs as well as to approximate the CCM volume. The dots tool of IMARIS was used to manually assign 3D coordinates (x,y,z) to the ECs within the experimental and control brain slices. The x,y,z positions of the ECs were exported for the nearest neighbor analysis.

### Image Analysis

The nearest neighbor (NN) distances were calculated using the knn.dist function of the Fast Nearest Neighbor (FNN) 1.1 R package. The computer simulations were performed with a random number generator in R followed by the same NN algorithm applied to the experimental and control brain slice data. All graphs were prepared with Prism 7 software. Error bars on applicable graphs are shown as mean + standard deviation.

### Statistical Analysis

All statistical analysis was performed with IBM SPSS Statistics Version 25. We tested the assumption of equal variance in our CCM and control NN distance data sets with Levene’s test. With the exception of the 2^nd^ NN pair for the CFP CCM and control data sets (Fig 3D), all data sets rejected the null hypothesis of equal variance; therefore, we used the Mann-Whitney test to compare CCM and control NN distances. Comparisons were considered statistically significant when p-value < 0.05.

## Results

### Visualization of somatically mutated endothelial cells in cerebral cavernous malformations with an inducible CCM3 mouse model and Confetti reporter

To study the development of cerebral cavernous malformations (CCMs) with respect to the endothelial cells (ECs) that have acquired the second, somatic mutation, we combined an endothelial-specific, tamoxifen-inducible cre recombinase (*PDGFb-iCreERT2*)^13^, lox-P flanked and knockout *Ccm3* alleles^14^, and the Brainbow2.1, hereafter referred to as confetti, reporter construct (*R26R-Confetti*)^15, 16^ in a transgenic mouse (Fig 1A). The experimental mice (*PDGFb-iCreERT2*, *Ccm3*^*fl/KO*^, *R26R-Confetti*^*fl/wt*^) provide a model in which a single, low dose of tamoxifen (TMX) results in simultaneous cre recombinase-mediated deletion of the single remaining floxed *Ccm3* allele and recombination within the *R26R-Confetti* allele to label mutant ECs and all of their progeny with a fluorescent signal. The confetti reporter is a widely utilized tool to randomly label cells with one of four different fluorescent proteins with varying cellular localizations: nuclear green fluorescent protein (nGFP), cytoplasmic red fluorescent protein (RFP), cytoplasmic yellow fluorescent protein (YFP), and membrane-bound cyan fluorescent protein (mCFP) (Fig 1B). We injected these transgenic mice with a single, 2-μg dose of TMX on either postnatal day 3, 4, or 5 to induce transient cre recombinase activity (Fig 1C). Following the single injection of TMX, a subset of endothelial cells within the vascular network acquired the induced second somatic mutation (deletion) and led to CCM development. CCM-null ECs overexpress phospho-myosin light chain (pMLC).^17, 18^ Due to the lack of a reliable antibody that recognizes Ccm3, and the desire to visualize a gain-of, rather than loss-of signal to mark the deletion of *Ccm3*, we utilized the increased phosphorylation of MLC as a secondary marker for *Ccm3* loss. Endothelial cells lining CCMs exhibit high levels of pMLC and are labeled with one of the confetti colors; thus, recombination at both the *Ccm3* and *R26R-Confetti* loci are observed in the same EC (Sup Fig. 1).

**Fig 1.**
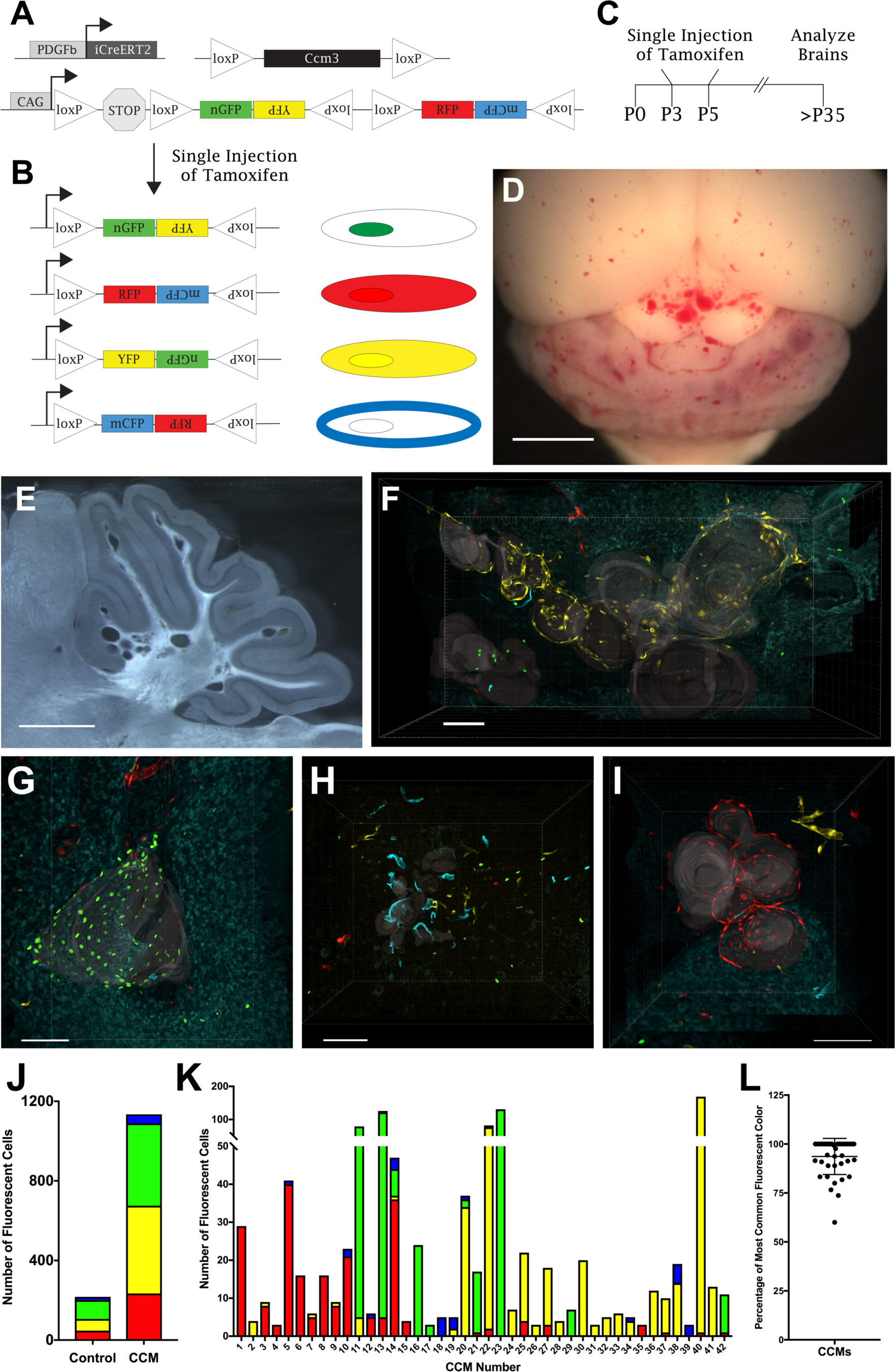
CCMs are composed of clonally dominant ECs labeled with a single confetti color. (A) Experimental mice contain three transgenes: endothelial-specific, tamoxifen-inducible cre recombinase with platelet derived growth factor b promoter (*PDGFb-iCreERT2*), loxP-flanked and null *Ccm3* alleles (*Ccm3*^*fl/KO*^), and *R26R-Confetti* reporter allele. (B) Cre recombination of a single *R26R-Confetti* allele leads to fluorescent labeling with one of four different constructs: nuclear GFP, cytoplasmic YFP, cytoplasmic RFP, and membrane-bound CFP. (C) Experimental design of a single, low dose (2μg) of tamoxifen injected into neonatal pups on postnatal days 3, 4, or 5. (D) *Ccm3*^*fl/KO*^, *PDGFb-iCreERT2*, *R26R-Confetti*^*fl/wt*^ experimental mice develop a modest CCM burden in the cerebellum and cerebral hemispheres (scale bar: 2mm). (E) A representative 80-μm thick brain slice with discrete CCMs dispersed throughout the cerebellum (scale bar: 1 mm). (F-I) 3D reconstructions of Z-series confocal images acquired across multiple serial brain slices to visualize the entire volume of CCMs in *Ccm3*^*fl/KO*^, *PDGFb-iCreERT2*, *R26R-Confetti*^*fl/wt*^ mice. Gray translucent surfaces have been added to the images to aid in visualizing the vascular lumens of each CCM. Representative CCMs for (F) YFP, (G) nGFP, (H) mCFP, and (I) RFP (scale bars: 100μm). (J) Number of ECs expressing each confetti color in serial brains slices of non-CCM control mice (*PDGFb-iCreERT2*, *R26R-Confetti*^*fl/wt*^, n=1) and CCM mice (*Ccm3*^*fl/KO*^, *PDGFb-iCreERT2*, *R26R-Confetti*^*fl/wt*^, n=3). (K) Number of each confetti color expressed by ECs lining individual CCMs (n=42). CCMs #40, #13, #18, and #s1,3-7,8 are the CCMs shown in panels F, G, H, and I respectively. (L) Percentage of the clonally dominant ECs, labeled with the most common confetti color within a CCM, when considering all confetti labeled ECs within the CCM (94+9 (mean + S.D.), n = 42).

Inducible CCM mouse models have been used extensively, with high doses and often multiple injections of TMX, to generate a severe phenotype to investigate the molecular signaling of CCM pathogenesis^19-25^ and efficacy of proposed therapeutics.^26-29^ Our approach was to more closely model the more severe familial form of the human disease, in which random second somatic mutations occur in a small population of endothelial cells, by using a very low dose of TMX. We induced a modest phenotype with CCMs dispersed throughout the cerebellum and cerebral hemispheres (Fig 1D). The dispersed CCMs enabled us to visualize individual CCMs to infer mutant EC lineage (Fig 1E). Serial brain sections, confocal microscopy, and 3D reconstruction were utilized to characterize the entire volume of individual CCMs. The Z-series of confocal images from the same brain region of serial slices were aligned and concatenated into 3D reconstructions of CCMs across large brain volumes. It was imperative to visualize an entire CCM, rather than a single section, to more confidently draw conclusions of the mutant EC lineage. Despite the use of a vibratome to section the entire brain into 80-μm thick slices, the sectioning process resulted in slight tissue loss between each slice, which is best demonstrated in a video of a rotating 3D reconstruction (Sup Video 1). The loss of tissue during brain processing was greatly outweighed by the valuable information gathered by reconstructing images from adjacent slices in order to examine entire CCMs.

### Clonal dominance of mutant ECs in CCMs

Representative CCM images for each confetti color demonstrate that CCMs are lined with a disproportionate number of ECs expressing the same confetti color (Fig 1F-I, Sup Vid 1-4). To measure the frequency of recombination within the *R26R-Confetti* allele with these experimental conditions in the absence of CCMs, we injected control animals (*PDGFb-iCreERT2*,*Ccm3*^*wt/wt*^, *R26R-Confetti*^*fl/wt*^) with the same dose of TMX on P3 (n=1). Serial brain slices of the P25 control mouse contained a fluorescent signal density of 1.3 confetti labeled ECs per 1×10^6^ mm^3^ of cerebellar brain volume. In the experimental brain slices, we occasionally observed CCMs containing a fluorescent EC density less than the 1.3 ECs per 1×10^6^ mm^3^ of brain volume that was measured as the background confetti labeling of the control sample. As every loxP-flanked allele, whether it is a gene trap or fluorescent reporter, has a unique recombination efficiency, there is a chance that during transient cre recombinase activity, recombination at two or more different alleles will not occur within the same cell. The CCMs that contained a confetti labeled EC density less than the control (non-lesion) samples are likely derived from mutant ECs in which the *R26R-Confetti* allele did not recombine to tag the mutant ECs with a color. This established the minimum value that could distinguish between chance recombination at the confetti locus and joint recombination of the *Ccm3* and *R26R-Confetti* alleles. Therefore, any CCMs with a fluorescent EC density less than 1.3 ECs per 1×10^6^ mm^3^ were not included in the analysis.

The CCM and control (non-lesion) samples had approximately equal proportions of ECs expressing each of the confetti colors (Fig 1J). The frequency of the *R26R-Confetti* allele recombining to tag ECs with each of the confetti colors within CCMs was: Red – 20.6%, Yellow – 38.9%, Green – 36.5%, and Cyan – 4.0%. This low rate of the *R26R-Confetti* allele tagging ECs with mCFP provided a rare event to more confidently conclude that multiple ECs expressing mCFP within a CCM are derived from a single mutant EC. Furthermore, the ability to detect mCFP in all samples enabled us to confidently conclude that there was no cre recombination event within the confetti locus in the unlabeled ECs. As detailed later, this ability to delineate ECs which have undergone cre recombination, from those ECs which have not, is important in our analysis of CCM development. While the experimental and control samples expressed each of the confetti colors in comparable frequencies, there was an overrepresentation of a single confetti color in labeled ECs within individual CCMs. To gain an accurate view of the molecular events occurring with the developing CCM, the number of ECs labeled with each of the confetti colors within 42 individual CCMs, of varying sizes and stages of maturation, was counted (Fig 1K). The clonal dominance of ECs tagged with the same confetti color was calculated, as the percentage of ECs labeled with the clonally dominant color when considering all ECs tagged with a confetti color within the CCM, to be 94%+9% (n=42) (Fig 1L). The disproportionate number of ECs tagged with the same confetti color in these malformations suggests clonal expansion of a mutant EC during CCM development.

### Nearest neighbor analysis validates the clonal dominance of mutant ECs in CCMs

Although even a cursory calculation of the difference in percentage of ECs labeled with each confetti color within individual CCMs showed a strong clonal dominance of one color (somatic mutation), we sought a more rigorous statistical analysis of the data. We performed a nearest neighbor (NN) analysis of labeled ECs using CCM lesion, control (non-lesion) tissue, and computer-generated simulation datasets. The NN algorithm is a method for determining the 3D distance between each member of a dataset. From the perspective of a single data point, the NN algorithm calculates the distance to all of the other data points and then ranks the distances from closest to furthest. The algorithm continues to the next data point within the group until all of the NN distances have been calculated and ranked for each member of the dataset. We applied the NN algorithm to the 3D reconstructions of the entire CCMs gathered from confocal images of serial brain slices. We represented each EC labeled with the clonally dominant confetti color within a CCM as a point with x,y,z coordinates (Fig 2A,B). The NN distances were calculated and ranked for a single EC in the CCM (visually depicted in Fig 2C, Sup Vid 5) and plotted as distance (μm) vs. NN number (Fig 2I). The NN algorithm continued to a second EC in the CCM (visually depicted in Fig 2D, Sup Vid 5), until the NN distances for all of the ECs within the CCM were calculated. The NN distances for all of the ECs within this representative CCM are plotted (Fig 2J).

**Fig 2.**
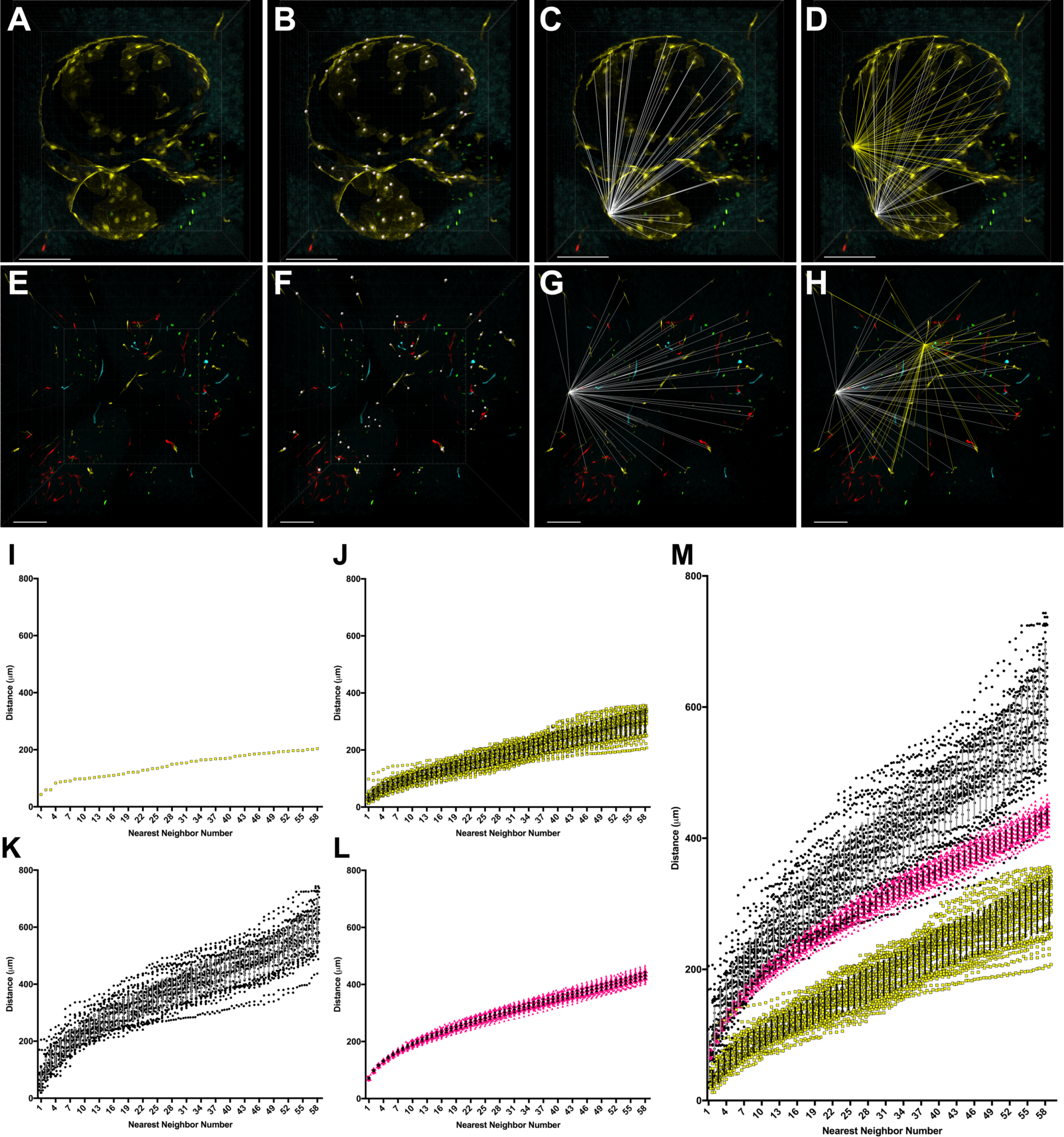
Nearest neighbor (NN) analysis as a quantitative comparison of the 3D distribution of confetti labeled ECs in CCMs, control brain slices, and computer simulations. (A) 3D reconstruction of the entire volume of a CCM containing clonally dominant ECs labeled with YFP. This CCM corresponds to CCM #22 in Figure 1K. (B) Representation of each EC labeled with YFP within the CCM as a point with x,y,z coordinates. (C) Visual representation of the 3D distances (white) between one EC and all of the other ECs within the CCM. The NN algorithm calculates and ranks these distances from closest to furthest. (D) Visual representation of the 3D distances (yellow) between a second EC and all of the other ECs within the CCM superimposed on the 3D distances of the first EC (white). (E-H) The same NN analysis is applied to non-CCM control brain slices: (E) 3D reconstruction of the control brain slices, (F) representation of each YFP labeled EC with x,y,z coordinates, (G) NN distances (white) between one YFP EC and all of the other YFP ECs, (H) a second set of NN distances (yellow) for a different YFP EC and all of the YFP ECs within the sample. (I-M) The NN distances visually represented on the 3D reconstructions above are ranked from closest to furthest and plotted as distance (μm) by NN number. (I) The NN distances between a single YFP EC within the CCM and all of the other YFPs within the CCM. This plot corresponds to the distances visually depicted in panel C above. (J) The NN distances for all of the YFP ECs within the CCM shown in panel A above. (K) The NN distances for all of the YFP ECs within the control brain slices shown in panel E above. (L) The average NN distances from 100 computer simulations in which the x,y,z coordinates of each data point were assigned by a random number generator. The simulation used the same volume and number of data points (ECs) as the experimental CCM sample shown in panel A. (M) The NN distances for the CCM (yellow), control brain slices (black), and computer simulations (pink) (error bars represent mean + standard deviation). All scale bars are 100μm.

The same NN algorithm was applied to serial brain slices of the control, *PDGFb-iCreERT2*, *Ccm3*^*wt/wt*^, *R26R-Confetti*^*fl/wt*^, mouse (Fig 2E-H, Sup Vid 6). These animals develop no CCM lesions and provide data on the background level of confetti recombination in otherwise identical conditions of transient cre recombinase induction. ECs labeled with the same confetti color within the control brain slices were each represented with an x,y,z point and the NN distances were calculated for comparison with the NN distances calculated for the CCM. Given the varying recombination frequencies of the four confetti colors (Fig 1J), each color was analyzed independently in both the CCM and control samples. Thus, the NN distances for YFP labeled ECs within the control brain slices are plotted, since the representative CCM being analyzed contained clonally dominant ECs expressing YFP (Fig 2K). Imaging experiments have the potential for experimenter bias to be introduced when selecting a region of tissue to study. For example, a control region of low confetti labeling would bias in favor of the heavily-labeled CCMs. Thus, we deliberately imaged regions of the control brain that contained many ECs expressing all of the confetti colors within the field of view, but we also created a second control data set that was generated randomly by a computer program. Using the x, y, and z dimensions of the 3D reconstruction of the CCM images and the number of ECs labeled with the clonally dominant confetti color of the CCM, the computer code generated a control dataset by randomly assigning the same number of data points into the same 3D volume as the experimental sample. The NN analysis was performed with this randomly-generated dataset and then the code repeated the process utilizing the same input parameters for a total of 100 iterations. The average NN distances for 100 simulations are plotted (Fig 2L).

Combining the NN analysis for a representative CCM, non-lesional control samples, and a computer-generated random simulation, the stark difference in the NN distances between the control and experimental groups strongly supports the hypothesis that the observed close proximity of the clonally dominant mutant ECs in the CCM is not due to random chance (Fig 2M).

### Clonal dominance is observed in CCMs across all confetti colors

We performed the nearest neighbor (NN) analysis for all CCM and the control (non-lesion) samples; the NN distances are plotted as distance (μm) vs. nearest neighbor number for each of the four confetti colors (Fig 3A-D). All CCMs, regardless of the confetti color expressed by the ECs, contained a clonally dominant population of ECs whose close proximity in 3D was not observed in the control brain slices. The CCM and brain slice control NN distances were statistically different beginning with the 1^st^ NN for the RFP groups (p<0.001), YFP groups (p<0.001), nGFP groups (p<0.001), and mCFP groups (p<0.001) as determined by a Mann-Whitney test. These very low p-values are driven by the large number of NN distances involved in each comparison. As p-values do not comment on the biological process of an observation, the more important result is the clear divergence of the CCM and control NN distances as the NN number increases (Fig 3A-D). The clonally dominant ECs within each of the 42 individual CCMs analyzed were significantly closer in 3D space than their respective confetti color controls. This pattern of clonally dominant ECs was observed across different CCMs as well as different confetti colors and illuminates a fundamental process of clonal expansion of *Ccm3*-null endothelial cells in CCM development.

**Fig 3.**
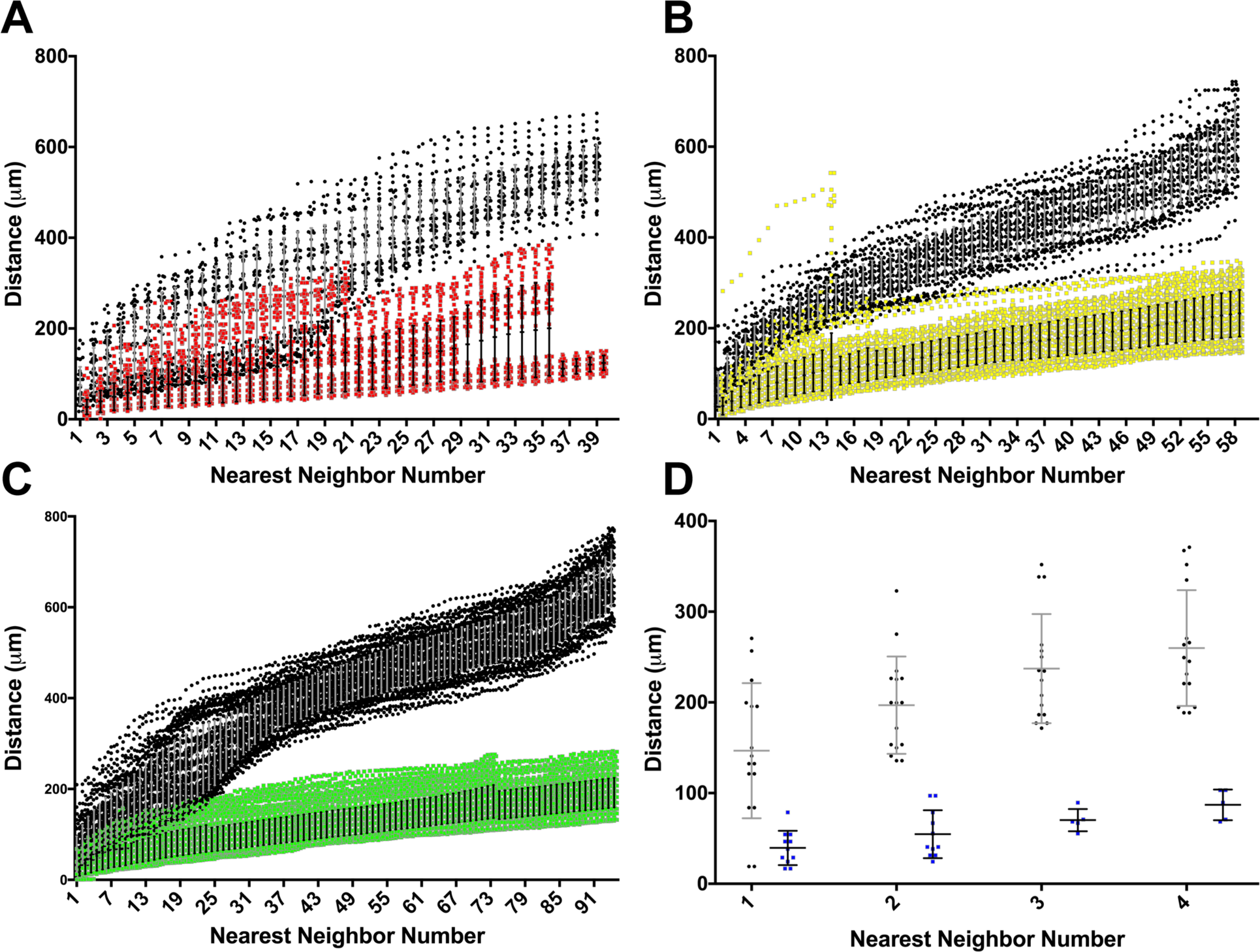
Divergence of the nearest neighbor (NN) distances of CCMs and non-lesion controls as NN number increases. Given the difference in recombination frequency among the four confetti colors, the NN comparison is only made between the ECs in CCMs and control brain slices labeled with the same confetti color. (A) NN distances for all CCMs containing clonally dominant ECs labeled with RFP (red) and the NN distances of ECs labeled with RFP in the control brain slices (black) (CCM n=13). (B) NN distances for YFP ECs within CCMs (yellow) and control brain slices (blank) (CCM, n=18). (C) NN distances for nGFP ECs within CCMs (green) and control brain slices (blank) (CCM, n=8). (D) NN distances for mCFP ECs within CCMs (blue) and control brain slices (blank) (CCM, n=3). For each of the four confetti colors, the NN distance was greater for the non-lesion controls than for the CCMs for every NN number pair, beginning with the 1^st^ NN (Mann-Whitney test, p<0.001).

### Clonal expansion of mutant ECs and recruitment of neighboring ECs drives CCM development

The increased number of confetti labeled ECs lining CCMs, that is not observed in the control samples, supports a clonal expansion of mutant ECs during CCM formation. Upon closer inspection of the CCMs imaged, a difference in the EC composition of small (diameter < 100μm) and large (diameter > 100μm) CCMs emerged. Many of the small CCMs imaged were composed nearly entirely of ECs labeled with the same confetti color (Fig 4A). Small CCMs consisting almost exclusively of clonally dominant ECs suggests that the EC which acquired the second somatic mutation, and was labeled with a confetti color, proliferated to initiate malformation development. We analyzed a small CCM with multiple caverns, some visibly connected within a single brain slice, in which each cavern was lined with ECs tagged with RFP (Fig 4A). Interestingly, these serial brain slice images also suggest that multicavernous CCMs can develop from the same somatic mutation. A rotation of the 3D reconstruction of the concatenated confocal images acquired across several serial brain slices more clearly demonstrates the high proportion or RFP-labeled ECs lining this multicavernous CCM (Sup Vid 7). In a different mouse, we visualized a larger multicavernous CCM (largest diameter > 200 μm) that contained both confetti-labeled and unlabeled ECs (Fig 4B). The unlabeled ECs appear as blank spaces within the vascular walls for the malformation. These unlabeled ECs within the CCM likely represent wildtype ECs that have been incorporated into the growing malformation. The possibility of these unlabeled ECs being *Ccm3*-null, having undergone cre recombination within the *Ccm3* allele and not the *R26R-Confetti* allele, cannot be excluded. However, as demonstrated in the pMLC staining of CCMs (Sup Fig 1) CCMs contain a mosaicism of mutant and wildtype ECs. The mosaicism of mutant and wildtype ECs has been previously shown with staining of human CCMs^30^ as well as human CCM DNA sequencing studies in which a small percentage of the total DNA reads detected the second somatic mutation of the mutant ECs within the CCM.^9-12^ In large CCMs we observe the same clonal expansion of mutant ECs as was seen with the small CCMs as well as the recruitment of unlabeled, presumably wildtype, neighboring ECs into the growing malformation. Taken together, we propose a model of CCM pathogenesis by which an EC acquires a second somatic mutation, undergoes clonal expansion through a gain in proliferative capacity, and recruits neighboring ECs as the size and complexity of the malformation increases (Fig 4C).

## Discussion

Herein we report a microscopy-based mutant EC lineage study of cerebral cavernous malformations in an inducible mouse model. Utilizing the confetti reporter, serial brain sectioning, confocal microscopy, and 3D reconstructions, we visualized entire CCMs and investigated CCM development at the cellular level. We studied both single-cavern and multicavernous CCMs of varying sizes that contained clonally dominant mutant ECs labeled with the same confetti color. The nearest neighbor analysis comparing CCMs, with non-CCM control brain slices, and a computer simulation determined that the large quantity of ECs labeled with the same confetti color in close 3D space that was seen in the CCMs could not be attributed to background confetti labeling during normal vascular development or random chance. These findings suggest that a single second somatic mutation is sufficient to seed the formation of a CCM. The final important observation from these studies was the incorporation of unlabeled, wildtype ECs into the growing CCMs. It appears that CCMs increase in size by incorporating wildtype ECs into the vessel wall of the malformation. We propose a model of CCM development by which an EC acquires a second somatic mutation, undergoes clonal expansion, and recruits neighboring ECs to expand the malformation.

**Fig 4.**
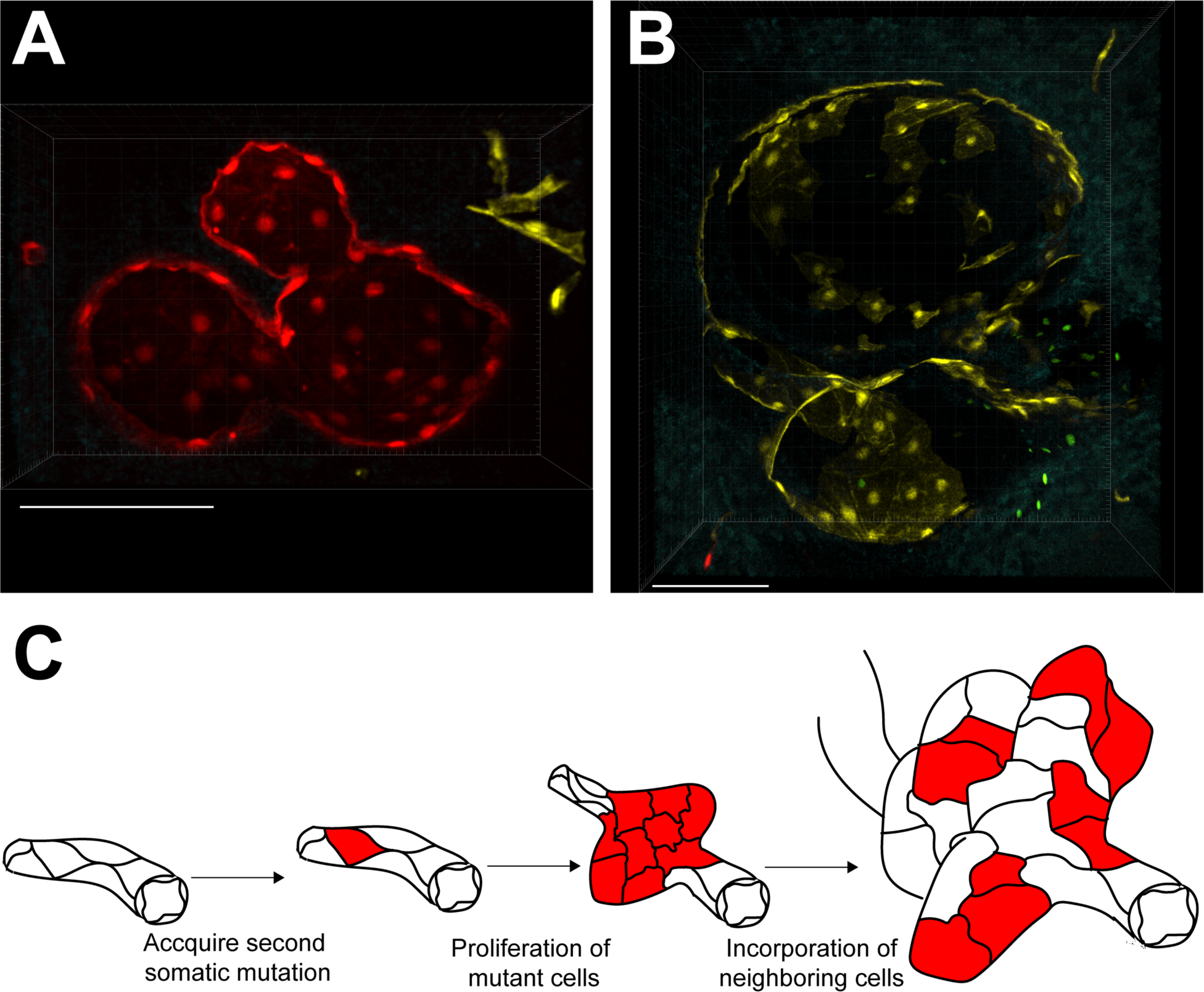
Clonal expansion of mutant ECs and recruitment of neighboring ECs drives CCM development. (A) Small (diameter <100μm), multicavernous CCM composed, nearly entirely, of clonally dominant ECs labeled with the same confetti color (RFP) in each cavern (scale bar: 100μm). This CCM corresponds to CCM #5 in Figure 1K. (B) Large (diameter >100μm), multicavernous CCM with a clonally dominant population of ECs labeled with YFP interspersed with unlabeled (appearing as blank spaces) ECs (scale bar: 100μm). This CCM corresponds to CCM #22 in Figure 1K. (C) Working model of CCM development in which a single, second somatic mutation leads to clonal expansion and subsequent recruitment of wildtype ECs into growing CCMs.

These results are consistent with and build upon the findings of several other research groups. Increased proliferation has been observed with astrocytes harvested from this inducible *Ccm3* mouse model and maintained in cell culture for several days.^14^ We published a DNA sequencing study of human CCMs in which one of the cases identified the somatic mutations of a surgically-resected CCM as well as a subsequent CCM that occurred later in the same anatomic location, presumably due to regrowth of a remnant of the original CCM. We found that both the initial and second CCM harbored the identical *CCM3* somatic mutation.^11, 12^ The formation of a second CCM from the same population of mutant ECs strongly supports our model of clonal expansion of a mutant EC in CCM development. These DNA sequencing studies also concluded that the frequency at which the somatic mutation was detected was less than 10% of all the DNA sequencing reads.^12^ The low detection rate of the second somatic mutation from large human CCMs highlights the mosaicism of CCM-null and wildtype ECs. This mosaicism with CCMs was first demonstrated with immunohistochemistry of human CCMs.^30^ Our work has expanded this concept of EC mosaicism by utilizing confocal microscopy to trace mutant ECs, with single-cell resolution, and observe the change in EC composition as CCMs develop. A pattern of CCM development emerged in which many of the small, recently formed CCMs consisted nearly entirely of clonally expanded mutant ECs while the large, multicavernous CCMs consisted of both the clonally expanded mutant ECs as well as a second population of unlabeled ECs. These two stages of CCM development: clonal expansion and incorporation of wildtype ECs may offer new opportunities for therapeutic intervention.

While CCMs are not considered vascular tumors, the increase in proliferation and regrowth of a malformation suggests that CCM-null ECs acquire an aggressive phenotype which may be targeted with anti-proliferative therapies. Work by others has targeted cell proliferation in the context of angiogenesis, with anti-angiopoietin-2 antibodies, and observed a reduction in CCM development in this neonatal CCM3 mouse model.^24^ Additional studies of anti-proliferative therapies, particularly those not directly targeting angiogenesis, may reveal a fundamental roles of CCM3 in both physiologic and pathologic conditions. Although the presence of wildtype ECs in CCMs has been inferred by the low mutant allele frequency observed in CCM tissue, the mechanism by which these ECs become incorporated in the growing malformation is completely unknown. As our understanding of this phenomena improves, we can use CCM mouse models like that of the current study to test therapeutics administered to disrupt the cellular signals involved in CCM incorporation of wildtype ECs.

In conclusion, this work is the first to provide visual evidence for clonal expansion of mutant ECs in the development of CCMs. By imaging the entire CCM volume with single-cell resolution, we captured the initial clonal burst of mutant ECs in small CCMs followed by the incorporation of wildtype ECs into the network of clonally dominant mutant ECs of larger CCMs. This work has illuminated the mutant EC lineage of CCMs and highlighted the need to further investigate the molecular mechanisms by which a single somatic mutation, within a single EC, leads to a large, functional vascular malformation that contains a mixed population of both mutant and wildtype cells.

## Acknowledgments

We would like to thank Amy Pieper, Heidy Pardo, Erin Griffin, Carol Gallione, and Christian Benavides for animal husbandry, Drs. Lisa Cameron and Benjamin Carlson of the Duke University Light Microscopy Core Facility for microscopy assistance, Dr. Ravi Karra of Duke University for helpful discussions, and Drs. Mark Kahn (University of Pennsylvania), Issam Awad (University of Chicago), and Mark Ginsberg (University of California at San Diego) for helpful discussions.

## Online Supplement

**Sup Fig 1.**
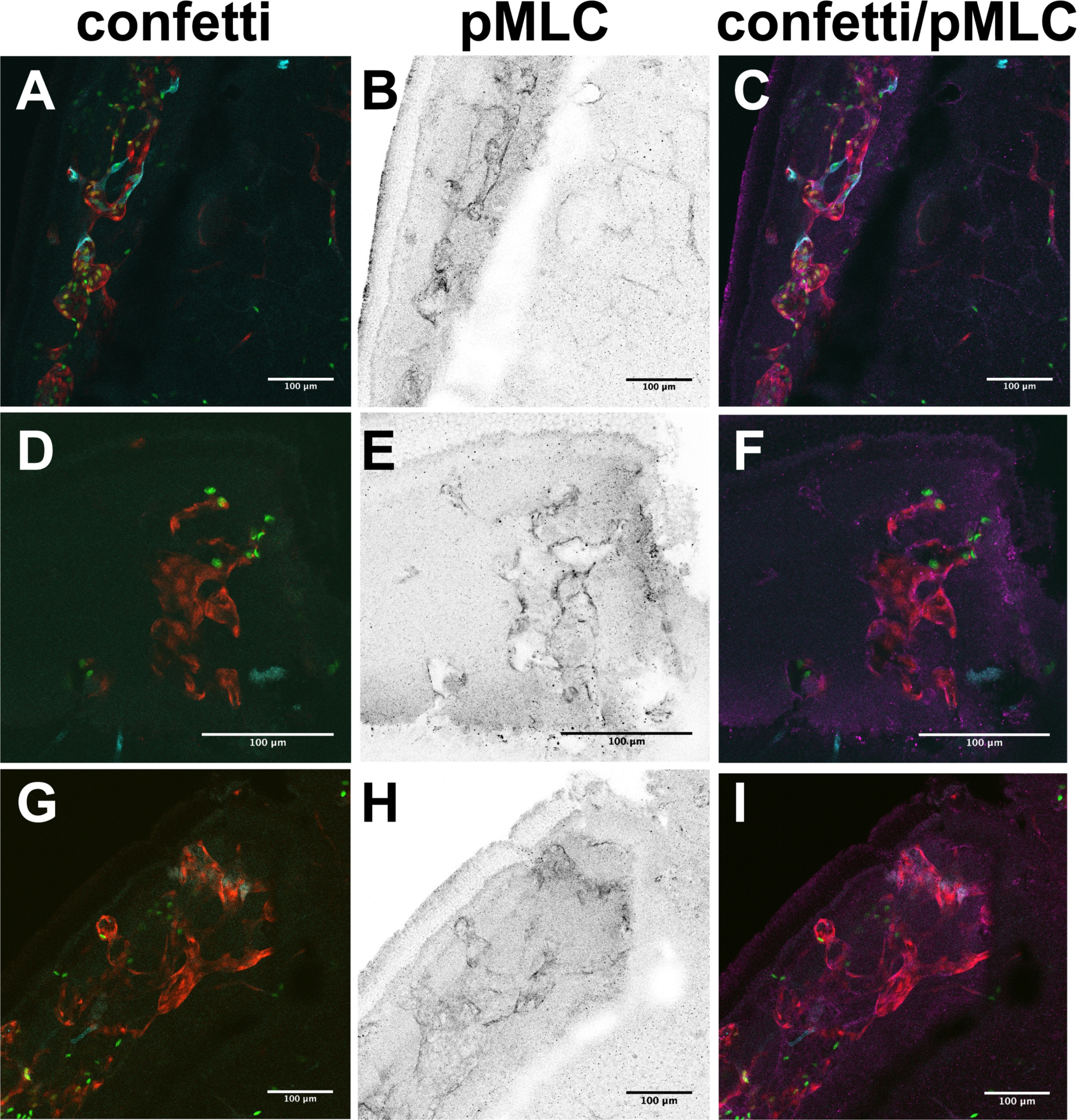
Transient cre recombinase activity deletes the *Ccm3* allele and recombines the *R26R-Confetti* allele within individual ECs that line CCMs. The developing vasculature at the periphery of the retina becomes malformed in this CCM mouse model and provides a tractable tissue to perform whole mount antibody staining. To increase the malformation burden, these mice (*Ccm3*^*fl/KO*^, *PDGFb-iCreERT2*, *R26R-Confetti*^*fl/wt*^) were injected with 25μg of tamoxifen. Phospho-myosin light chain (pMLC) is upregulated in Ccm-null cells and was used as a secondary marker for *Ccm3* deletion.^1,2^ The colocalization of a confetti color and the increase in pMLC signal identifies an individual EC in which the *Ccm3* and *R26R-Confetti* alleles have undergone cre recombination. The confetti colors, pMLC staining with Alexa Fluor 647, and the composite image are shown for three regions of CCMs (A-C, D-F, G-I). No YFP labeled ECs were detected in these regions. The lack of signal is likely due to YFP denaturation during the heat-induced antigen retrieval necessary for the pMLC staining. The ECs within these malformations that are labeled with a confetti color also have an increase in pMLC staining. Thus, both recombination events (deletion of the *Ccm3* allele and labeling with a confetti color) are able to occur within the same EC of this mouse model.

**Sup Video 1. 3D reconstruction of 6 serial brain slices containing a large, multicavernous CCM composed of clonally dominant ECs labeled with YFP.** Gray translucent surfaces have been added to this 3D reconstruction to aid in visualizing the CCM lumens. This sample also contains a second CCM, located in the lower left-hand region, composed of clonally dominant ECs labeled with nGFP. All four confetti colors are visualized throughout this sample. The ability to detect each confetti color reduces the probability that an unlabeled EC (appearing as a blank region of the CCM) has undergone cre recombination as we are able to detect all ECs which have recombined at the confetti locus. The background level of confetti labeling in the brain regions that are not part of the CCM also demonstrate the low rate of cre recombination we induced with the single, 2-μg dose of tamoxifen.

**Sup Video 2. 3D reconstruction of 2 serial brain slices containing a large, single-cavern CCM composed of clonally dominant ECs labeled with nGFP.** Gray translucent surfaces have been added to this 3D reconstruction to aid in visualizing the CCM lumens. This sample also contains other CCMs, located in the upper region, composed of clonally dominant ECs labeled with RFP.

**Sup Video 3. 3D reconstruction of 4 serial brain slices containing a small CCM composed of clonally dominant ECs labeled with mCFP.** Gray translucent surfaces have been added to this 3D reconstruction to aid in visualizing the CCM lumens.

**Sup Video 4. 3D reconstruction of 4 serial brain slices containing multiple small, single-cavern and multicavernous CCMs composed of clonally dominant ECs labeled with RFP.** Gray translucent surfaces have been added to this 3D reconstruction to aid in visualizing the CCM lumens.

**Sup Video 5. Visual representation of the NN distances for 2 YFP ECs within a large CCM composed of clonally dominant ECs labeled with YFP.** The first rotation is the 3D reconstruction of the CCM used for the sample NN analysis described in Figure 2. The second rotation visually depicts the distances from one EC to all of the other ECs within the CCM. The third rotation visually depicts the distances from a different EC to all of the other ECs within the CCM. While not shown in the video, the NN algorithm continues until the distances between every EC within the CCM have been calculated and ranked from smallest to largest. The complete set of NN distances are shown in Figure 2J.

**Sup Video 6. Visual representation of the NN distances for 2 YFP ECs within the non-lesion control brain slices.** The first rotation is the 3D reconstruction of the control brain slices used for the sample NN analysis described in Figure 2. The second rotation visually depicts the distances from one YFP-labeled EC to all of the other YFP-labeled ECs within the sample. The third rotation visually depicts the distances from a different YFP-labeled EC to all of the other YFP-labeled ECs within the sample. While not shown in the video, the NN algorithm continues until the distances between every YFP-labeled EC within the control brain sample have been calculated and ranked from smallest to largest. The complete set of NN distances are shown in Figure 2K.

**Sup Video 7. 3D reconstruction of a small, multicavernous CCM composed of clonally dominant ECs labeled with RFP.** Gray translucent surfaces have been added to this 3D reconstruction to aid in visualizing the CCM lumens. This 3D reconstruction contains the CCM shown in Fig 4A.

**Sup Video 8. 3D reconstruction of a large, multicavernous CCM composed of clonally dominant ECs labeled with YFP along with unlabeled, wildtype ECs.** Gray translucent surfaces have been added to this 3D reconstruction to aid in visualizing the CCM lumens. This 3D reconstruction contains the CCM shown in Fig 4B as well as a second CCM containing clonally dominant ECs labeled with nGFP.

